# Dual Inhibition of sEH and COX-2 Improved Cognition in Alzheimer’s Disease via Enhanced Myogenic Response and Cerebral Artery Distensibility

**DOI:** 10.1101/2025.08.21.669945

**Authors:** Gilbert C. Morgan, Andrew Gregory, Chengyun Tang, Sung Hee Hwang, Jane J. Border, Jing Xu, Yedan Liu, Shan Bai, Tae Jin Lee, Cameron Cantwell, David Bunn, Karen M. Wagner, Christophe Morisseau, Carly Pittman, Alina Ngo, Peter Osayi, Aditi Pabbidi, Philip O’Herron, Zsolt Bagi, Jessica A. Filosa, Hongwei Yu, Cindy McReynolds, Bruce D. Hammock, Richard J. Roman, Fan Fan

## Abstract

Genetic studies have linked *EPHX2* (encoding soluble epoxide hydrolase, sEH) and *PTGS2* (encoding cyclooxygenase-2, COX-2) to Alzheimer’s disease (AD). Elevated levels of sEH and COX-2 found in AD patients and animals suggest their involvement in neurodegeneration, glial activation, vascular dysfunction, and inflammation. This study evaluated the effects of a new dual sEH/COX-2 inhibitor, PTUPB, on cerebrovascular function and cognition in TgF344-AD rats. The rats received oral PTUPB (2 mg/kg/day) for 25 days. Body weight, plasma glucose, and HbA1c levels remained stable between PTUPB- and vehicle-treated AD rats. PTUPB significantly improved recognition memory in AD rats, as detected by the Novel Object Recognition test. Pressure myography showed that PTUPB restored myogenic responses and increased the distensibility of the middle cerebral arteries (MCAs) in AD rats. Acute PTUPB (0.1 and 1 μM) enhanced myogenic contraction in response to elevated perfusion pressure in AD MCAs, with minimal effects in wild-type vessels. Vehicle-treated AD rats displayed impaired functional hyperemia, whereas PTUPB (1 μM) significantly restored this response. Transcriptomic analysis of cerebral vascular smooth muscle cells from AD rats indicated that PTUPB influences genes related to contractility, extracellular matrix remodeling, inflammation, and oxidative stress. These results provide new evidence that dual inhibition of sEH and COX-2 improves cognition in AD, likely by enhancing myogenic response and increasing cerebral artery distensibility. Our findings highlight the potential of PTUPB as a therapeutic approach for cerebrovascular dysfunction in AD.

## INTRODUCTION

Soluble epoxide hydrolase (sEH) and cyclooxygenase-2 (COX-2) are critical enzymes involved in distinct branches of arachidonic acid (AA) metabolism, which contribute to the regulation of inflammation, vascular function, and homeostasis.^1, 2^ COX-2 converts AA into prostaglandins (PGs), which are lipid mediators that play an essential role in pain, fever, inflammation, vascular reactivity, platelet aggregation, and gastrointestinal protection.^3, 4^ In contrast, sEH metabolizes epoxy-eicosatrienoic acids (EETs) into dihydroxy-eicosatrienoic acids (DHETs). While EETs are AA-derived signaling molecules with anti-inflammatory, vasodilatory, and cardioprotective properties, DHETs may promote proinflammatory responses.^5^

Genetic studies have identified *EPHX2* (encoding sEH)^6–9^ and *PTGS2* (encoding COX-2)^10^ as genes linked to Alzheimer’s disease (AD). Both sEH^11–13^ and COX-2^14, 15^ are upregulated in AD patients and animal models. Inhibition or genetic deletion of either enzyme has been shown to improve cognitive function in AD and cerebral hypoperfusion animal models by modulating amyloid processing, tau phosphorylation, synaptic plasticity, inflammation, neuromodulation, metabolism, vascular regulation, and microglial function.^3, 10–13, 16–22^ These findings suggest that targeting sEH and COX-2 can help restore the disrupted AA signaling network in AD. By preserving EETs and reducing proinflammatory PGs, inhibition of these enzymes may shift the neuroinflammatory milieu toward a more protective state. In addition to its anti-inflammatory effects, sEH inhibition improves vascular reactivity and cerebral perfusion, alleviating chronic cerebral hypoperfusion, a key contributor to AD-related neurodegeneration.^12, 22–25^ Although direct evidence from AD models remains limited, recent studies in stroke^26^ and memory decline conditions^27^ demonstrate that COX-2 inhibition can restore vascular function by reducing blood–brain barrier (BBB) disruption and enhancing regional brain metabolism, the latter serving as a surrogate marker of improved cerebral perfusion and neurovascular coupling. Together, sEH and COX-2 inhibition may promote vascular homeostasis, reduce cerebrovascular dysfunction, and shift the neurovascular environment toward a more neuroprotective state, through either additive or synergistic effects.

We previously reported that chronic sEH inhibition improved cerebral hemodynamics and cognition in AD and diabetes-related Alzheimer’s Disease-Related Dementias (ADRD) animal models. ^6, 12, 25^ The highly selective sEH inhibitor 1-(1-propanoylpiperidin-4-yl)-3-[4-(trifluoromethoxy)phenyl]urea (TPPU) improved cognitive function and cerebrovascular health in AD and diabetes-related ADRD models. These beneficial effects were associated with reduced oxidative stress, amyloid burden, and hippocampal neuron loss; mitigation of capillary rarefaction and glial activation; enhanced neurovascular coupling and BBB integrity; and increased expression of both pre- and post-synaptic proteins. Sun et al. also reported that TPPU alone offers neuroprotective and anti-neuroinflammatory effects in AD mouse models.^21^ Recent studies have demonstrated that PTUPB, a dual sEH/COX-2 inhibitor,^28^ exhibits broad therapeutic potential across multiple preclinical models, including alleviation of LPS-induced acute lung injury,^29^ suppression of tumor growth,^30^ reduction of inflammatory pain^31^ and nephrotoxicity.^32^ Importantly, by inhibiting sEH, PTUPB can increase EET levels, potentially counteracting the negative cardiovascular effects of COX-2 inhibition by maintaining a more favorable balance between prostacyclin and thromboxane.^30, 31^ PTUPB has also been shown to mitigate blood pressure elevation and proteinuria, which are associated with cardiovascular risk.^32^

Despite the evidence supporting selective sEH inhibition in AD/ADRD, most studies on PTUPB have focused on non-neurological conditions. To date, no studies have explored the effects of dual sEH/COX-2 inhibition in AD or ADRD. The present study aims to address this gap by assessing the impact of PTUPB on cerebrovascular function and cognitive performance in a TgF344-AD rat model of AD.

## MATERIALS AND METHODS

### General

TgF344-AD (AD) rats and age-matched Fischer 344 (F344) controls were used for experiments. PTUPB, provided by Dr. Hammock’s laboratory, was dissolved in 100% polyethylene glycol, molecular weight 400 (PEG-400; 91893, MilliporeSigma, Burlington, MA) and subsequently diluted in drinking water to achieve a final PEG-400 concentration of 1%. Wild-type (WT) Fischer 344 rats *(F344/NHsd)* were obtained from ENVIGO (Indianapolis, IN), and TgF344-AD rats *[F344-Tg(Prp-APP, Prp-PS1)19]* were sourced from the Rat Resource & Research Center (RRRC) at the University of Missouri (Columbia, MO), then bred and housed at Augusta University under a 12-hour light-dark cycle with *ad libitum* food and water. Beginning at 26 weeks of age, F344 and AD rats were treated for 25 days with either vehicle (1% PEG-400 in drinking water) or PTUPB (2 mg/kg/day in the vehicle). Genotyping was performed by PCR to confirm the presence of the APP and PS1 transgenes in TgF344-AD rats, using primers and protocols provided by the RRRC. All animal protocols were approved by the Institutional Animal Care and Use Committees (IACUC) of Augusta University and were conducted in accordance with NIH guidelines for the care and use of laboratory animals.

### Body Weight, Plasma Glucose, and Glycated Hemoglobin A1c (HbA1c)

Body weight, plasma glucose, and HbA1c levels were assessed before and after chronic PTUPB treatment in conscious rats, approximately 3 hours after the start of the lights-on cycle. Blood samples were collected from the tail vein, as previously described. ^25, 33, 34^ Plasma glucose was measured using a Contour Next glucometer (Ascensia Diabetes Care, Parsippany, NJ), and HbA1c levels were measured using an A1CNow+ device (Polymer Technology System, Indianapolis, IN).

### Pressure Myography

#### MCA Isolation

MCAs were isolated based on our previously established protocols.^35–39^ Animals were humanely euthanized under 4% isoflurane anesthesia, and brains were rapidly harvested, weighed, and immersed in ice-cold calcium-free physiological salt solution (PSS_0Ca_), composed of 119 mM NaCl, 18 mM NaHCO₃, 10 mM glucose, 5 mM HEPES, 4.7 mM KCl, 1.18 mM NaH₂PO₄, 1.17 mM MgSO₄, and 0.03 mM EDTA (pH 7.4). A 5 × 3 mm cortical section containing the MCA was carefully dissected and transferred to PSS_0Ca_ supplemented with 1% bovine serum albumin. The pia mater was gently separated, and branch-free M2 segments of the MCA (100–200 μm in diameter) were identified and isolated under a stereomicroscope, then kept in ice-cold PSS_0Ca_ for further analysis.

#### Myogenic Response of the MCA

Branch-free M2 segments of the MCA were carefully positioned in a pressure myograph chamber (Living Systems Instrumentation, Burlington, VT) by securing both ends onto glass micropipettes (1B120-6, World Precision Instruments, Sarasota, FL). The pressure myograph chamber was maintained at 37 °C and continuously perfused with a calcium-containing physiological salt solution (PSS_Ca_) composed of 1.6 mM CaCl₂ and aerated with a gas mixture of 21% O₂, 5% CO₂, and 74% N₂. The chamber was positioned on an IMT-2 inverted microscope (Olympus, Center Valley, PA) equipped with a digital imaging system (MU1000, AmScope, Irvine, CA). Following cannulation, the vessels were stretched to their approximate *in situ* length and pressurized to 40 mmHg. They were then allowed to equilibrate for 30 minutes to permit the development of spontaneous tone.^40–42^ Vessels were considered viable if they exhibited a vasoconstrictive response greater than 15% when exposed to 60 mM KCl in PSS_Ca_.^43^ Those failing to meet this threshold were excluded from further analysis. After equilibration and confirmation of viability, the inner diameters (IDs) of MCAs were recorded from rats that had received either vehicle or PTUPB treatment for 25 days. To evaluate the myogenic response, pressure–diameter relationships were measured across a range of transmural pressures, from 40 to 180 mmHg, in 20 mmHg increments, following our previously established protocol.^44, 45^

#### Myogenic Tone of the MCA

To assess passive vessel diameter, the intraluminal pressure was readjusted to 40 mmHg, and the arteries were rinsed 5–6 times with PSS_0Ca_ to eliminate extracellular calcium. Under these conditions, passive inner diameters (ID₀_Ca_) were recorded across the same pressure range used during active tone assessment. Myogenic tone was then calculated using the formula:

Myogenic tone (%) = [(ID_0Ca_ – ID) / ID_0Ca_] X 100, following our previously described protocol.^39, 46, 47^

#### Cross-sectional area (CSA) of the MCA

CSA (*μ*m^2^) was calculated using the formula:

CSA = (*π*/4) X (OD^2^_0Ca_ − ID^2^_0Ca_), where OD_0ca_ and ID_0Ca_ represent the outer and inner diameters of the MCA measured in PSS_0Ca_, respectively.^39, 46, 47^

#### Distensibility of the MCA

Distensibility (%) was calculated using the formula:

Distensibility = [(ID_0Ca_ − ID_0Ca5mmHg_) / ID_0Ca5mmHg_] X 100, where ID_0Ca5mmHg_ represents the ID measured at 5 mmHg in PSS_0Ca_.^39, 46, 47^

#### Incremental distensibility of the MCA

Incremental distensibility (% / mmHg) was calculated using the formula:

Incremental distensibility = [ΔID_0Ca_ / (ID_0Ca_ X ΔP)] X 100, where P represents perfusion pressure, ΔID_0Ca_ is the change in inner diameter between two perfusion pressures measured under calcium-free conditions (PSS₀_Ca_), ID₀Ca is the inner diameter at current pressure, and ΔP is the difference in perfusion pressure (mmHg).^39, 46, 47^

#### Acute PTUPB Treatment on Myogenic Response of the MCA

These experiments were performed using TgF344-AD and F344 control rats aged 27–30 weeks. MCAs were freshly isolated and cannulated with glass micropipettes in a pressure myograph chamber, as per our previously established protocols.^44, 45^ Vessels were equilibrated for 30 minutes in PSS containing calcium to allow for the development of spontaneous myogenic tone. Vessel viability was confirmed by a robust constrictive response to 60 mM KCl in PSS_Ca_.^25, 40–42^ Following viability confirmation, the arteries were superfused with either vehicle (0.1% DMSO) or the dual sEH/COX-2 inhibitor PTUPB. For PTUPB-treated vessels, a cumulative concentration protocol was employed to assess dose-dependent effects on myogenic reactivity. PTUPB was applied sequentially at concentrations of 0.1, 1, and 10 µM (each prepared in 0.1% DMSO), starting with the lowest dose. After an initial equilibration period, pressure–diameter relationships were recorded over a range of transmural pressures (40 to 160 mmHg in 20 mmHg increments). The complete pressure-response curve was completed within approximately 30 minutes per dose to minimize time-dependent changes in vascular behavior. After data acquisition at each dose, vessels were thoroughly washed with fresh PSS_Ca_ and allowed to re-equilibrate at a baseline pressure of 40 mmHg before introducing the next higher concentration. This procedure ensured vessel viability and minimized the risk of residual drug effects or desensitization. To control for any time-dependent drift or mechanical fatigue, the response to 0.1% DMSO vehicle was tested separately on a paired vessel from the same animal using an identical protocol.

### Whisker Stimulation-Induced Functional Hyperemia

Whisker stimulation-induced functional hyperemia in response to PTUPB was evaluated in 27–30-week TgF344-AD and F344 control rats, following our established protocol.^12, 33, 34^ Briefly, rats were anesthetized using Inactin (100 mg/kg) and ketamine (60 mg/kg). Baseline arterial pressure was maintained within physiological levels (100– 120 mmHg), and end-tidal CO₂ was kept at 35 mmHg by adjusting the respiratory rate, monitored using an end-tidal CO₂ analyzer (CAPSTAR-100, CWE Inc., Ardmore, PA).^12, 33, 34, 39^ Using a low-speed air drill, a thinned-skull (semi-transparent) cranial window was carefully formed at a location 2 mm posterior and 6 mm lateral to the bregma. A laser speckle imaging system (RWD Life Science Co., Ltd., Shenzhen, China) was positioned approximately 10 cm above the window. To enhance image clarity, saline was applied to the surface, and the system was adjusted for CBF monitoring. To assess the CBF response to sensory stimulation, the right-side whiskers were stimulated at a frequency of 10 Hz for 1 minute, while CBF in the left somatosensory cortex was continuously recorded. This procedure was repeated in three separate trials at 5-minute intervals. To assess the dose-dependent effects of PTUPB on functional hyperemia, a stepwise concentration protocol was used. PTUPB (0.1, 1, and 10 μM in 0.1% DMSO, respectively) was sequentially applied to the thinned skull over the cranial window for 10 minutes prior to measuring the functional hyperemic response. Each dose-response measurement was then completed within approximately 10 minutes to limit time-dependent variability. Between doses, the skull surface was thoroughly rinsed with saline and allowed to return to baseline for 10 minutes before the next application. For vehicle controls, a single application of 0.1% DMSO was applied to a paired animal, and CBF responses to whisker stimulation were recorded using the same imaging setup.

### Bulk RNA-seq Analysis

Primary vascular smooth muscle cells (VSMCs) were isolated from the middle cerebral arteries of AD rats. The extraction procedure consisted of sequential enzymatic digestion, beginning with protease dithiothreitol (2 mg/mL) and papain (22.5 U/mL), followed by treatment with collagenase (250 U/mL), elastase (2.4 U/mL), and trypsin inhibitor (10,000 U/mL), in accordance with previously published protocols.^34, 38, 39, 46^ Cultures from three animals, at early passages (P2–P4), were maintained in Dulbecco’s Modified Eagle Medium (DMEM; Thermo Fisher Scientific, Waltham, MA) enriched with 10% fetal bovine serum (Thermo Scientific) and incubated at 37 °C in a 5% CO₂ atmosphere. Cells were exposed for 48 hours to either PTUPB at a final concentration of 1 µM or a vehicle control consisting of 0.1% DMSO. Total RNA from both control- and PTUPB-treated AD cerebral VSMCs was extracted using Trizol reagent (Thermo Fisher Scientific) and further purified with the PureLink RNA Mini Kit (Thermo Fisher Scientific). Assessment of RNA integrity, reverse-transcription quantitative PCR (RT-qPCR), preparation of sequencing libraries, and the RNA-seq runs were all performed at the Integrated Genomics Shared Resource, Georgia Cancer Center. Raw reads generated from high-throughput sequencing underwent adapter removal and quality trimming via Cutadapt v4.0.^48^ Cleaned sequences were mapped to the rat reference genome (rn7) using STAR v2.7.10a,^49^ and read counts for each gene were compiled. Differential expression analysis employed the DESeq2 package,^50^ and functional enrichment was conducted with clusterProfiler.^51^ For pathway analysis, annotations from the Kyoto Encyclopedia of Genes and Genomes (KEGG) database were used.

### Novel Object Recognition and Placement Tests

Two complementary behavioral assays were used to evaluate memory function in rats. <u>The novel object placement test</u> measures spatial recognition memory, which depends heavily on hippocampal function and also engages frontal cortex networks when assessing “episodic-like” recognition memory.^52, 53^ In this task, two identical objects were positioned in opposite corners of the chamber, and the rat was allowed to explore for 10 minutes during the acquisition phase. After a 1-hour delay, one object was moved to a new location, the rat was returned to the chamber, and exploration was recorded for 10 minutes. <u>The novel object recognition test</u> examines hippocampus-dependent non-spatial short-term memory.^52–54^ During the acquisition phase, two identical objects were placed in opposite corners of the arena, and each rat was allowed to explore for 10 minutes. After a 10-minute delay, one of the objects was replaced with a novel one, and the rat was returned for an additional 10-minute exploration period.

For both tasks, memory performance was quantified using:

Recognition Index (RI) = TN/(TN + TF)

Discrimination Index (DI) = (TN – TF)/(TN + TF), where TN = time spent exploring the novel object and TF = time spent exploring the familiar object.

RI represents the proportion of total exploration devoted to the novel stimulus. Positive DI values indicate a preference for the novel object/location, negative values indicate a preference for the familiar one, and a value of zero indicates no preference. A failure to preferentially explore the novel or relocated object was interpreted as a memory deficit, consistent with prior reports.^33, 55^ Between trials, the apparatus was cleaned thoroughly with 70% ethanol, followed by distilled water to eliminate odor cues.

### Statistics

All values are expressed as mean ± standard error of the mean (SEM). Group differences for longitudinal measures (body weight, plasma glucose, HbA1c) were assessed by two-way repeated-measures ANOVA (treatment × time), with Holm–Šidák *post hoc* test. Myogenic pressure–diameter curves and functional hyperemia dose– responses were analyzed by two-way repeated-measures ANOVA (treatment × pressure or dose) with Holm–Šidák *post hoc* test. Between-group comparisons used unpaired or paired two-tailed *t*-tests as appropriate, and one-way ANOVA with Tukey’s post *hoc test* for >2 groups. Behavioral index values were compared to chance performance (zero) using one-sample *t*-tests, and between-group differences were assessed by *t*-test or one-way ANOVA. Bulk RNA-seq differential expression was analyzed with DESeq2, with significance defined as false discovery rate (FDR)–adjusted *q* < 0.05. Functional enrichment analysis (clusterProfiler) was also performed using *q* < 0.05. All tests were two-tailed, and statistical analyses were performed using GraphPad Prism 10 (GraphPad Software) or R 4.3.1. Significance was set at *p* < 0.05.

## RESULTS

### Chronic PTUPB Treatment Did Not Alter Body Weight, Plasma Glucose, and HbA1c Levels in AD Rats

As presented in **Figure 1**, body weight increased modestly over time across all groups, with no statistically significant differences observed between treatments. On Day 0 and Day 25, body weights were as follows: F344 (Veh), 429.6 ± 13.9 g to 447.6 ± 13.4 g; AD (Veh), 450.6 ± 7.5 g to 471.0 ± 11.2 g; and AD (PTUPB), 459.4 ± 8.4 g to 466.4 ± 8.4 g. Plasma glucose and HbA1c levels showed no significant changes across the study period in any group, despite minor fluctuations being observed. All measurements were validated through diagnostic assessments, including Q-Q plots and residual plots, which confirmed the normality and homoscedasticity of residuals and supported the use of parametric statistical analyses.

**Figure 1.**
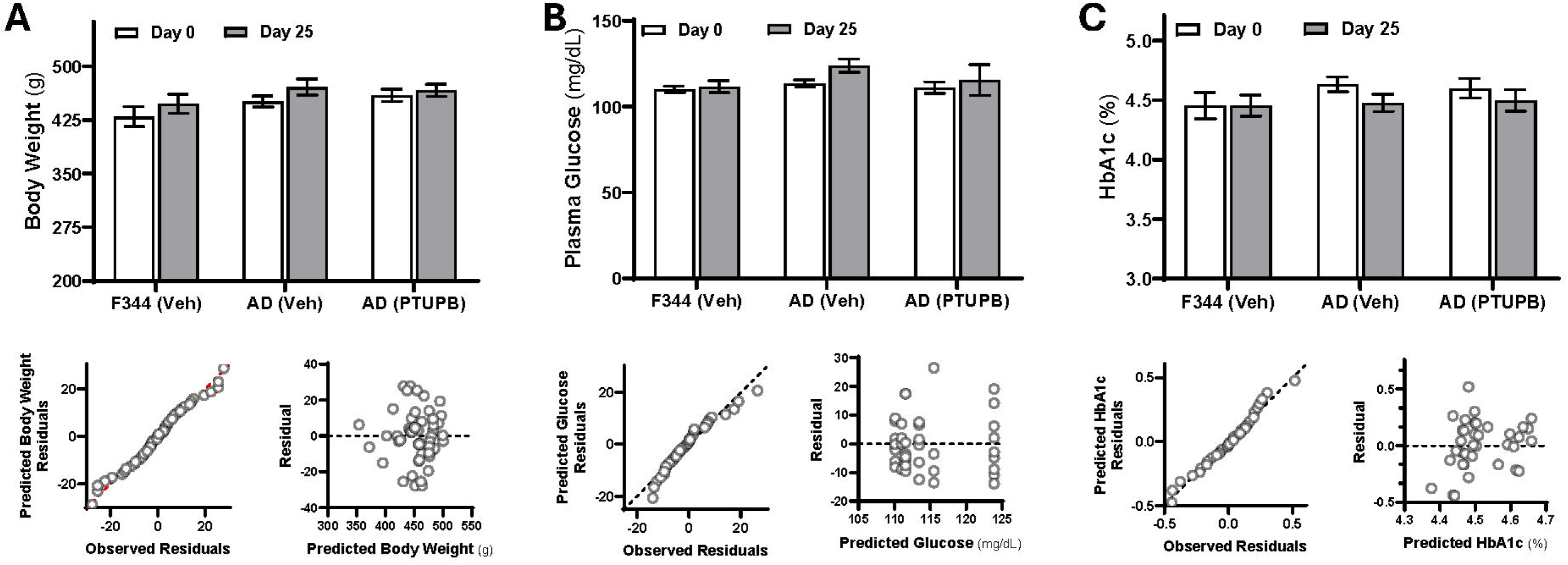
Chronic PTUPB Treatment Did Not Alter Body Weight, Plasma Glucose, and HbA1c Levels in AD Rats. Comparison of body weight (**A**), plasma glucose (**B**), and HbA1c (**C**) in age-matched vehicle (Veh)- and PTUPB-treated TgF344-AD (AD) vs. Veh-treated F344 (control) rats. Upper panels display group values. Lower left panels show Quantile-quantile (Q–Q) plots to assess normality of residuals, and lower right panels show residual plots to evaluate model assumptions. Mean values ± SEM are presented, with 4 - 9 rats per group.

### Chronic PTUPB Treatment Enhances Myogenic Responses and Vascular Distensibility of MCAs in AD Rats

Consistent with our earlier findings,^25, 45^ vehicle-treated AD rats demonstrated a significantly blunted myogenic response in the MCA compared to vehicle-treated F344 control rats (**Figure 2A**). In vehicle-treated control rats, the MCAs exhibited a robust myogenic response, constricting by 13.3 ± 1.9%, 16.8 ± 1.9%, and 5.6 ± 2.5% as perfusion pressure increased from 40 to 100, 140, and 180 mmHg, respectively. In contrast, AD rats showed a markedly impaired response, with vessels dilating by 0.92 ± 4.1%, 17.8 ± 5.5%, and 45.9 ± 7.6% at the same pressure levels, with forced dilation observed as early as 100 mmHg. However, in PTUPB-treated AD rats, this impairment was substantially reversed; MCAs constricted by 11.6 ± 2.7% at 100 mmHg and showed only modest dilation of 5.7 ± 3.7% and 20.6 ± 6.7% at 140 and 180 mmHg, respectively, indicating partial restoration of myogenic response.

**Figure 2.**
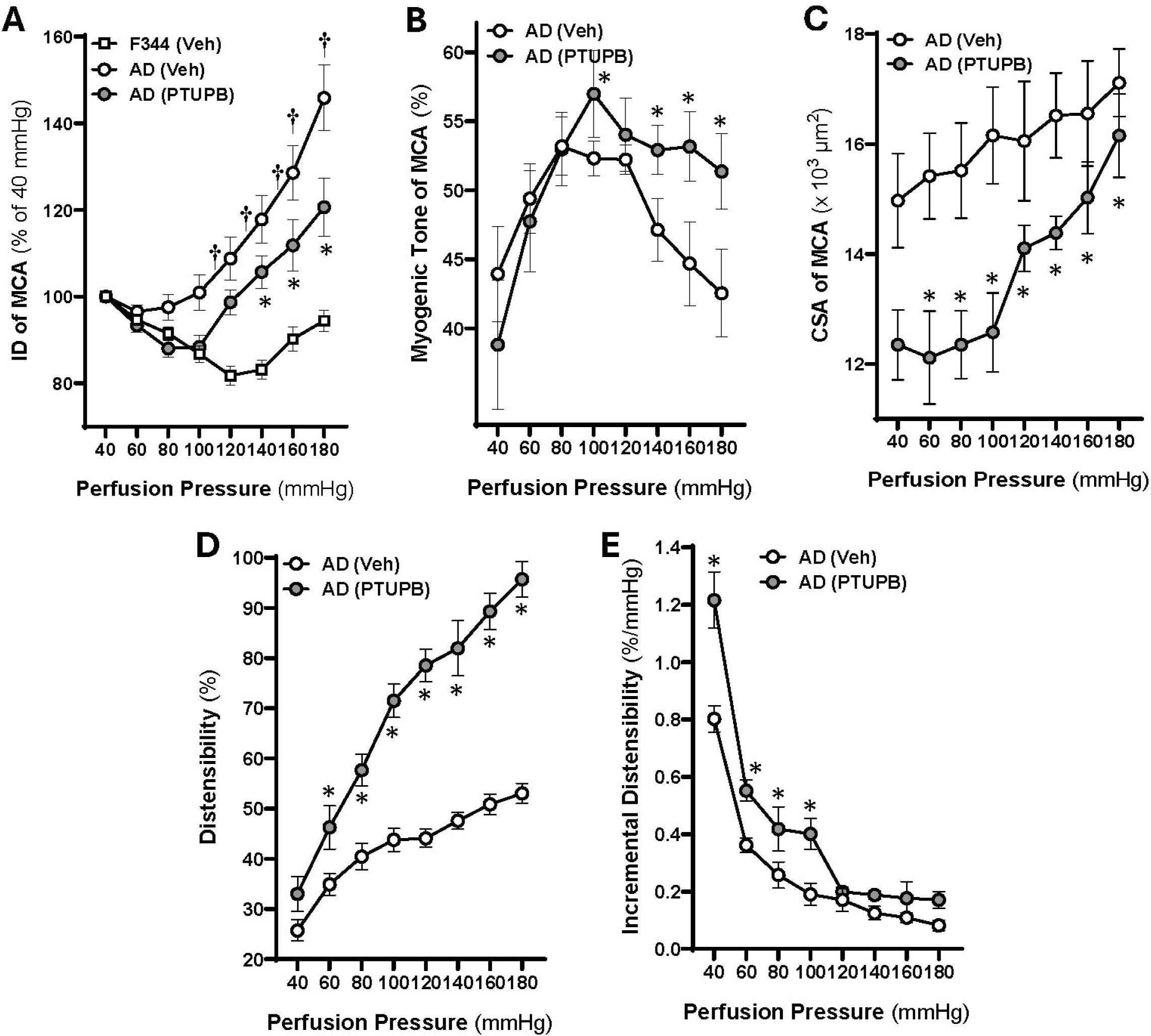
Chronic PTUPB Treatment Enhances Myogenic Responses and Vascular Distensibility of MCAs in AD Rats. Comparison of myogenic responses, tone, cross-sectional area (CSA), distensibility, and incremental distensibility in middle cerebral arteries (MCAs) from chronic vehicle (Veh)- and PTUPB-treated TgF344-AD (AD) and F344 (control) rats. **(A)** Myogenic response of the middle cerebral artery (MCA) in age-matched vehicle (Veh)- and PTUPB-treated TgF344-AD (AD) rats compared to Veh-treated F344 (control) rats. **(B–E)** Comparisons of myogenic tone **(B)**, CSA **(C)**, distensibility **(D)**, and incremental distensibility **(E)** of MCAs in Veh- and PTUPB-treated AD rats. Data are presented as mean ± SEM, with 10–20 vessels from 4–9 rats per group. **\*** *p* < 0.05 vs. Veh-treated AD rats; **†** *p* < 0.05 vs. Veh-treated F344 rats.

The effects of PTUPB treatment on myogenic tone, wall structure (CSA), and passive mechanical properties (distensibility and incremental distensibility) were also evaluated in AD rats compared to vehicle-treated AD controls. While PTUPB treatment did not significantly enhance myogenic tone at lower perfusion pressures, a significant increase was observed at 140 to 180 mmHg in vessels from PTUPB-treated AD rats compared to vehicle-treated AD rats. This finding further supports that PTUPB restores the impaired myogenic response in AD by enhancing pressure-dependent vasoconstriction (**Figure 2B**). Additionally, PTUPB treatment resulted in a significant reduction in CSA of MCA compared to vehicle-treated AD rats across most pressure levels (**Figure 2C**). Distensibility of the MCA was significantly increased from 60 to 180 mmHg in PTUPB-treated AD rats compared to vehicle-treated AD rats (**Figure 2D**). Furthermore, incremental distensibility was significantly elevated at perfusion pressures ranging from 40 to 100 mmHg in PTUPB-treated AD rats relative to vehicle-treated AD rats (**Figure 2E**).

### Acute PTUPB Treatment Enhances Myogenic Responses of the MCAs in AD Rats

As shown in **Figure 3**, the myogenic response of vehicle-treated MCAs in TgF344-AD rats was significantly impaired compared to vehicle-treated F344 vessels. In TgF344-AD rats, acute treatment with PTUPB at low doses (0.1 and 1 μM) significantly enhanced the myogenic response at higher pressures, whereas the 10 μM dose blunted this response. In contrast, PTUPB at low doses had no significant effect on the myogenic response in F344 vessels, while the 10 μM dose significantly attenuated the response. Interestingly, despite the initial differences in baseline myogenic function, there were no significant differences in the pressure–diameter relationships between PTUPB-treated F344 and TgF344-AD vessels at any given dose, suggesting that PTUPB treatment normalized myogenic behavior in the AD model.

**Figure 3.**
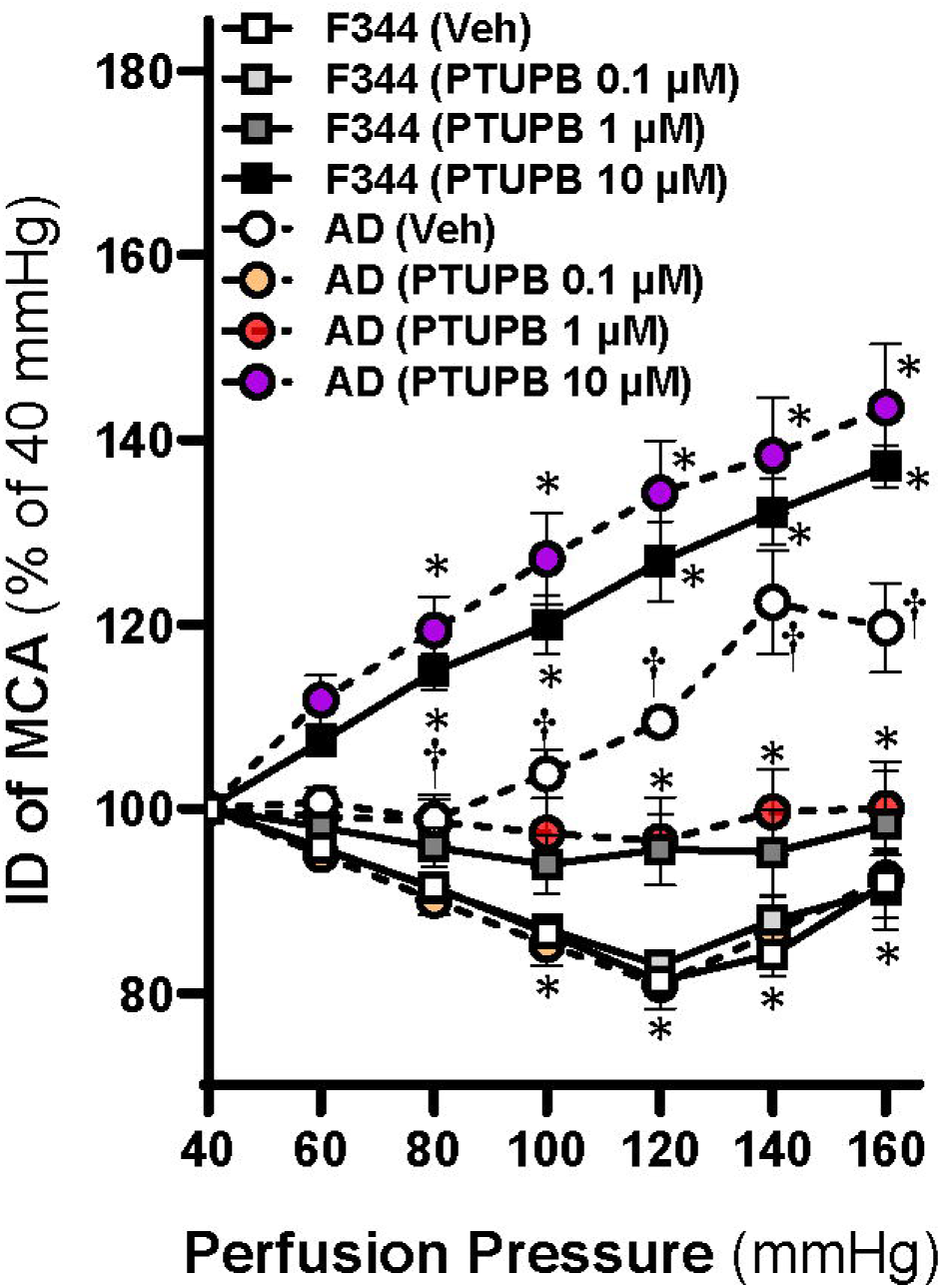
Acute PTUPB Treatment Enhances Myogenic Responses of the MCAs in AD Rats. Comparison of myogenic responses in middle cerebral arteries (MCAs) from acute vehicle (Veh)- and PTUPB-treated TgF344-AD (AD) and F344 (control) rats. Data are presented as mean ± SEM, with 6-16 vessels from 4–8 rats per group. **\*** *p* < 0.05 vs. Veh- treated rats; **†** *p* < 0.05 vs. F344 rats at the corresponding drug and dose

### PTUPB Restores Whisker Stimulation-Induced Functional Hyperemia in AD Rats

**Figure 4A** compares whisker stimulation–evoked functional hyperemia in response to PTUPB between 27–30-week-old TgF344-AD rats and age-matched F344 controls. At all tested concentrations, the CBF increase was significantly attenuated in AD rats compared with F344 controls. PTUPB at 1 μM markedly enhanced functional hyperemia in both strains relative to vehicle-treated rats within the same strain (**Figure 4B**).

**Figure 4.**
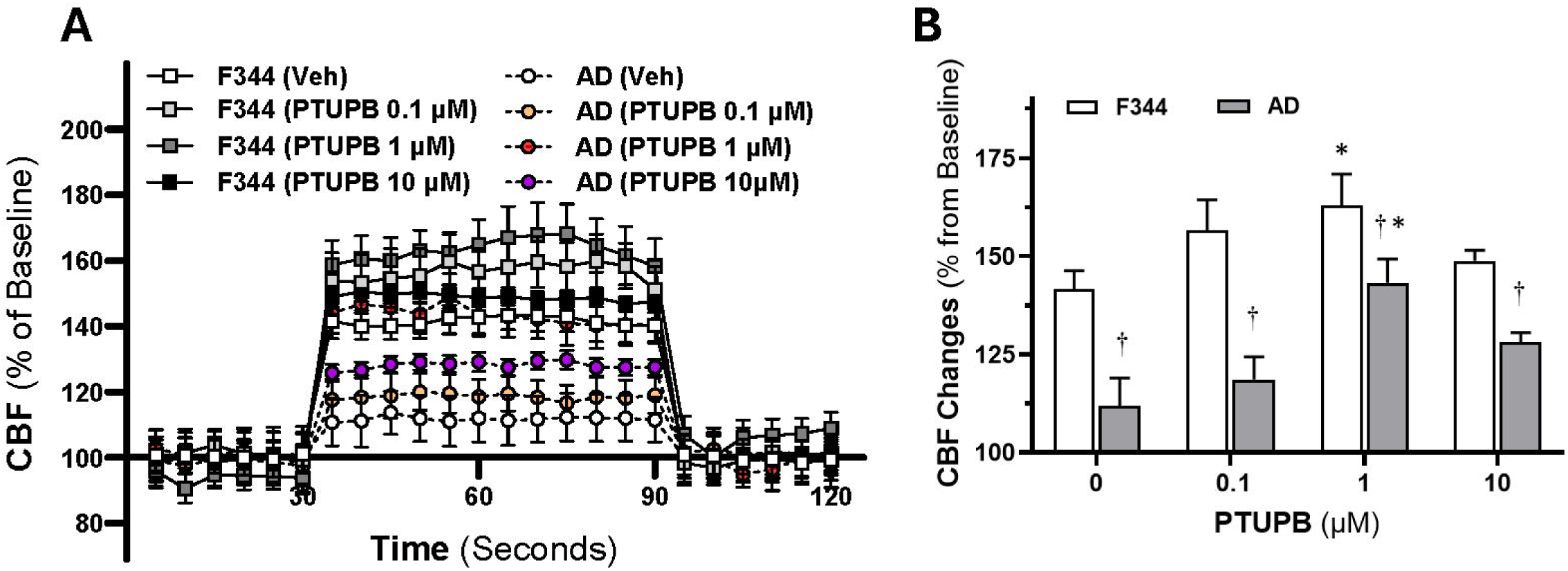
Acute PTUPB Treatment Restores Whisker Stimulation-Induced Functional Hyperemia in AD Rats. Comparison of whisker stimulation–evoked functional hyperemia in response to PTUPB between TgF344-AD rats and age-matched F344 controls. **(A)** Time course of CBF changes during whisker stimulation (30–90 s). **(B)** Mean CBF changes over the stimulation period (30–90 s). Data are presented as mean ± SEM (4–8 rats per group). **\*** *p* < 0.05 vs. Veh-treated rats; **†** *p* < 0.05 vs. F344 rats at the corresponding drug and dose.

### PTUPB Modulates Pathways Involved in Contractile Function, Extracellular Matrix Remodeling, Inflammation, and Oxidative Stress in AD VSMCs

Bulk RNA-seq analysis of primary VSMCs from AD rats demonstrated significant modulation of multiple signaling pathways following PTUPB treatment (**Figure 5A**). The most enriched included focal adhesion, ECM–receptor interaction, PI3K–Akt, and TGF-beta signaling. Additional enriched pathways comprised VSMC contraction, regulation of the actin cytoskeleton, Rap1 signaling, and adherents junction. Pathways related to inflammation and oxidative stress were also altered, including MAPK, AGE–RAGE, TNF signaling, fluid shear stress and atherosclerosis, mTOR, and HIF-1 signaling. Oxytocin signaling was significantly enriched. The volcano plot (**Figure 5B**) depicts transcriptomic differences between PTUPB-treated and vehicle-treated AD VSMCs, highlighting numerous significantly upregulated (red) and downregulated (blue) genes.

**Figure 5.**
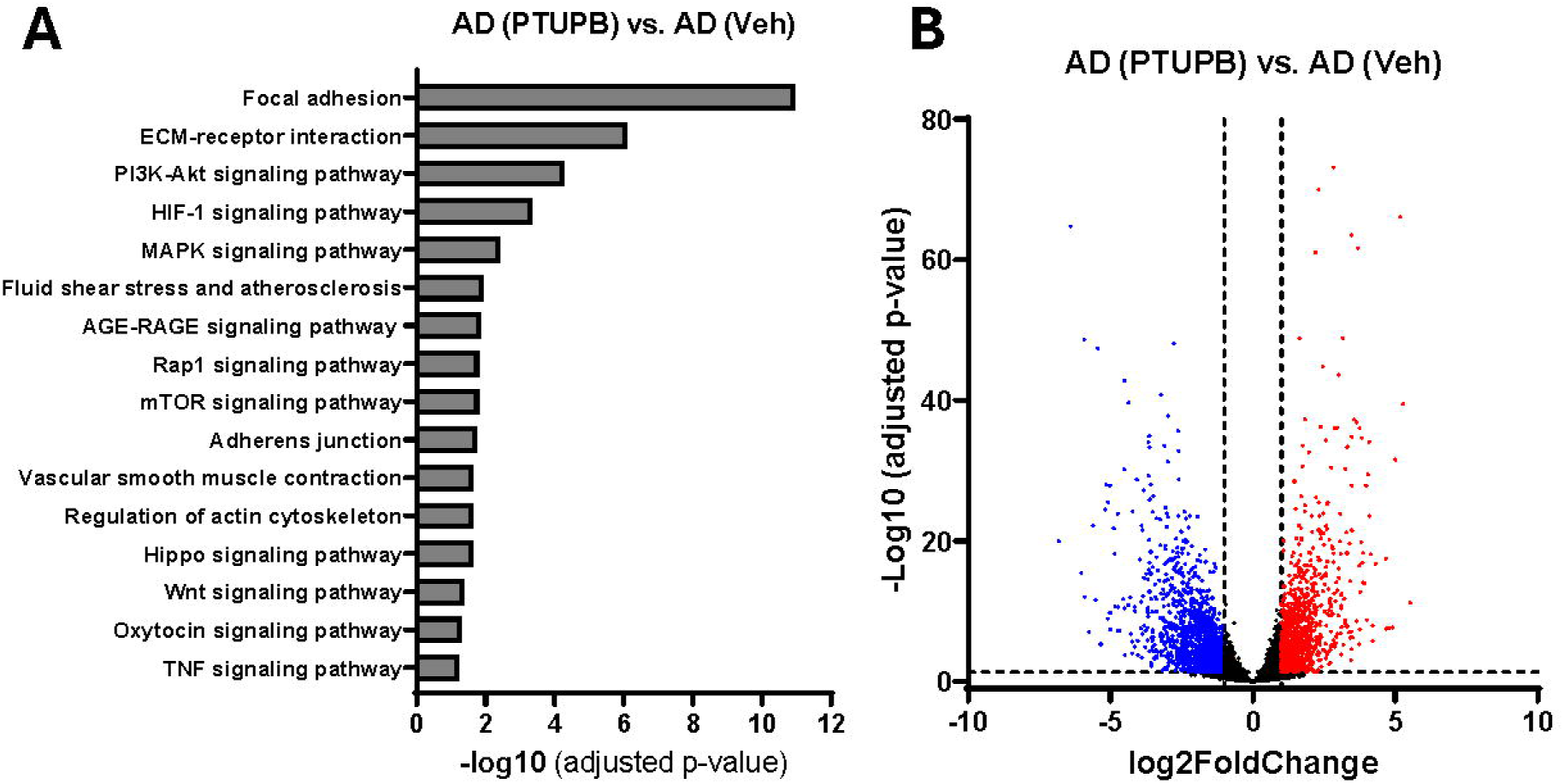
PTUPB Modulates Pathways Involved in Contractile Function, Extracellular Matrix Remodeling, Inflammation, and Oxidative Stress in AD VSMCs. Comparison of transcriptomic profiling by bulk RNA-seq in primary cerebral vascular smooth muscle cells (VSMCs) from Veh- and PTUPB-treated AD rats. **(A)** KEGG-based enrichment analysis highlighting the most significantly impacted pathways following PTUPB exposure in AD VSMCs (FDR-adjusted *p* < 0.05). **(B)** Volcano plot illustrates gene expression differences between PTUPB and veh-treated AD rats. The horizontal axis depicts the log2 fold change, and the vertical axis shows the negative log10 *p*-value. Genes meeting the thresholds of |log2FC| > 1 and an FDR-adjusted *p* < 0.05 (or *q*-value) are marked in red for upregulation and blue for downregulation.

### PTUPB Restores Memory Performance in Novel Object Recognition and Placement Tests in AD Rats

In the novel object recognition test (**Figure 6A**), vehicle-treated AD rats showed a significantly lower recognition index than F344 controls, indicating impaired non-spatial memory, consistent with our previous reports.^25, 45^ PTUPB treatment restored recognition index values in AD rats to near-control levels. In the novel object placement test (**Figure 6B**), vehicle-treated AD rats had a markedly reduced discrimination index, consistent with spatial memory deficits. PTUPB treatment significantly improved discrimination index values, restoring a preference for the novel location.

**Figure 6.**
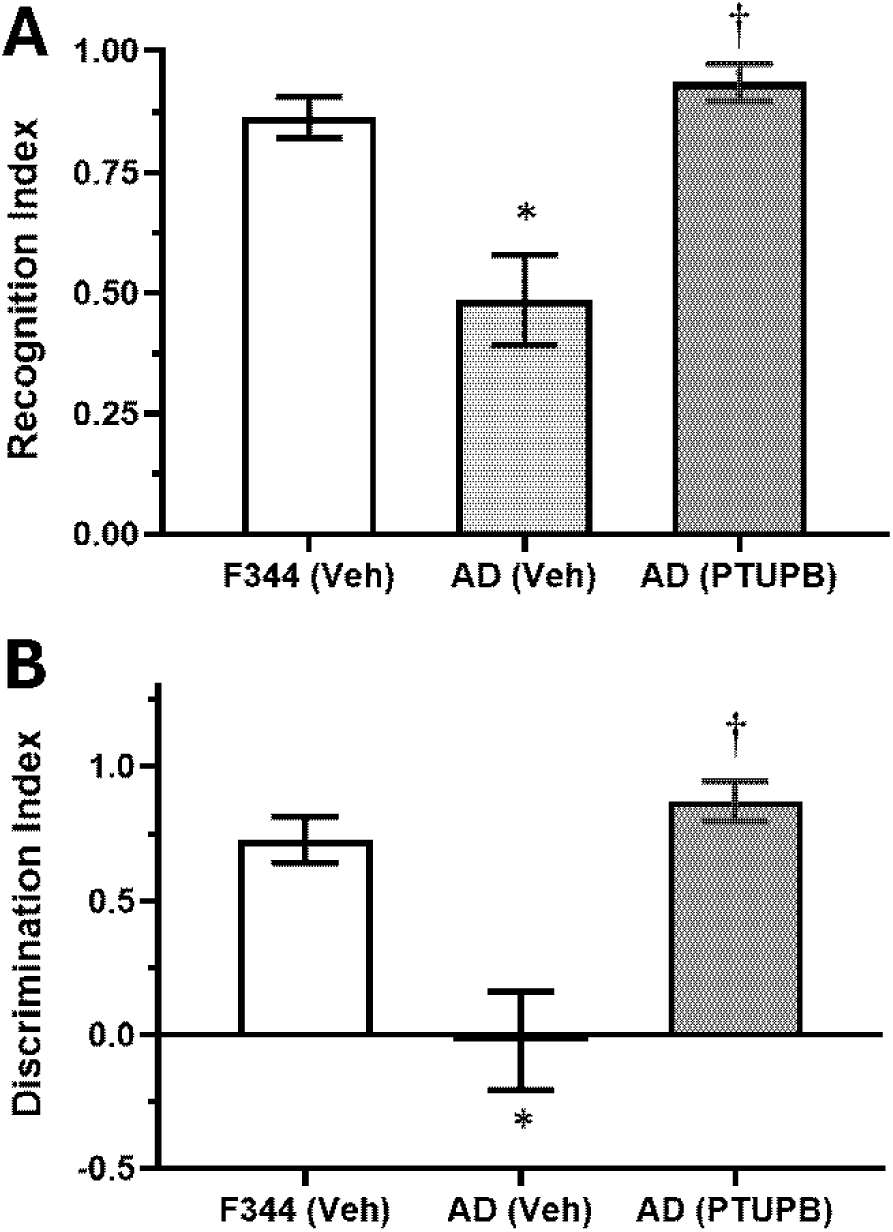
PTUPB Restores Memory Performance in Novel Object Recognition and Placement Tests in AD Rats. Performance in behavioral assays comparing vehicle-treated F344 controls, vehicle-treated TgF344-AD rats, and PTUPB-treated TgF344-AD rats. (**A**) Recognition index in the novel object recognition test, assessing non-spatial short-term memory. (**B**) Discrimination index in the novel object placement test, assessing spatial recognition memory. Data are mean ± SEM (5–12 rats per group). **\*** *p* < 0.05 vs. F344 (Veh); **†** *p* < 0.05 vs. AD (Veh).

## DISCUSSION

AD is a progressive, degenerative brain disorder and the predominant cause of dementia across the globe. Although the full disease mechanism is not yet completely understood, the prevailing amyloid-beta (Aβ) cascade hypothesis suggests that an abnormal buildup of Aβ initiates a series of harmful events, including tau protein hyperphosphorylation, formation of neurofibrillary tangles, neuronal loss, and eventual cognitive decline.^56^ In recent years, therapeutic development has shifted away from managing symptoms toward eliminating Aβ deposits. Anti-Aβ monoclonal antibodies such as lecanemab and donanemab have shown the ability to reduce plaque burden and modestly slow disease progression. However, treatment with these agents is frequently complicated by amyloid-related imaging abnormalities (ARIA).^57^ In donanemab clinical studies, ARIA-E (brain swelling or fluid accumulation) was reported in roughly 20–24% of participants, and ARIA-H (microhemorrhages or superficial siderosis) in about 27–31%. According to FDA labeling, 36.8% of individuals receiving donanemab experienced some form of ARIA, compared with 14.9% in the placebo group. People carrying the APOE ε4 allele, especially those with two copies, face an even greater risk because of increased vascular fragility, inflammatory responses, and the disruptive effects of rapid plaque clearance.^58^ These observations highlight the possibility that concentrating solely on the Aβ cascade may overlook other important factors. Considering vascular injury, genetic influences, and other overlapping disease mechanisms may sharpen our understanding of AD and guide the design of safer, more effective treatments.

Vascular injury in AD is increasingly recognized as a central contributor to disturbances in CBF, and brain hypoperfusion is a common finding in AD and other dementia-related disorders.^59, 60^ In the brain, blood flow is maintained by finely tuned regulatory processes, including autoregulation and neurovascular coupling.^61, 62^ Autoregulation stabilizes CBF to protect delicate capillaries from fluctuations in arterial pressure. Myogenic responses are the major contributors to CBF autoregulation. Failure of this system increases pressure stress on capillaries, inducing BBB leakage, glial activation, neurodegeneration, and cognitive impairment.^61, 63–65^ Impaired autoregulation is strongly associated with cerebral small vessel disease, stroke, AD, hypertension, and diabetes-related dementia.^33, 64, 66–68^ Notably, maintaining arterial pressure above the lower autoregulatory limit during surgery has been shown to reduce postoperative delirium and enhance memory recovery.^69^ In parallel, neurovascular uncoupling, where blood flow responses no longer match neuronal activity, also contributes to hypoperfusion. Under physiological conditions, increased local neuronal activity induces functional hyperemia, which is mediated in part by Kir2.1 channel activation in capillary endothelia.^70, 71^ In AD, this coupling is disrupted, worsening energy deficits and driving neurodegeneration.^33, 71, 72^

Vascular inflammation is a potent amplifier of these vascular injuries. In AD/ADRD, inflammatory mediators and immune cell activation disrupt the BBB, diminish CBF autoregulation, and exacerbate hypoperfusion and neuronal loss. COX-2 drives the production of pro-inflammatory prostanoids,^4^ while sEH degrades anti-inflammatory, vasoprotective EETs into more inflammatory DHETs.^6–9^ Genetic evidence links *PTGS2* (encoding COX-2) and *EPHX2* (encoding sEH) to AD/ADRD, and both enzymes are elevated in patient samples and experimental models, with effects spanning amyloid processing, tau pathology, synaptic function, immune activation, vascular tone, and microglial activity.^11–15^ While COX-2 or sEH inhibition individually improves cognition in animal models, each strategy carries limitations. The present study evaluated a promising approach using PTUPB, a dual COX-2/sEH inhibitor developed by EicOsis,^28^ which may enhance anti-inflammatory efficacy while reducing cardiovascular toxicity. In earlier work, we found that sEH inhibition with TPPU reduced astrocyte activation, decreased hippocampal amyloid plaques, restored MCA myogenic response, improved CBF autoregulation, lowered vascular oxidative stress, and improved cognition in AD and DM-ADRD rats.^12, 25^

The TgF344-AD rat model overexpresses mutated human amyloid precursor protein (APP) and presenilin 1 (PS1).^73^ This AD model exhibits Aβ plaque, gliosis, and learning dysfunction from 6 months.^73^ Prior studies, including our own, indicate BBB breakdown, neurofibrillary tangles, neurodegeneration, and impaired CBF response to hypercapnia or pressure in this model. ^25, 45, 70, 74, 75^ In the current study, we evaluated the effects of PTUPB on cognition and cerebral vascular function in TgF344-AD rats at an age when cognitive impairment is well established.^45, 73^ Since we previously reported that sEH inhibition reduced plasma glucose and HbA1c levels in a diabetes-related ADRD model, ^25^ we first assessed body weight, glucose, and HbA1c levels in the present study. We found no significant differences in these parameters between PTUPB- and vehicle-treated AD rats or their controls. We found that PTUPB restored both recognition memory and spatial memory, indicating improvement in hippocampal- and cortical-dependent cognitive function. This cognitive benefit was associated with an enhanced myogenic response of the MCA. Moreover, PTUPB increased myogenic tone in the range of 140– 180 mmHg in AD rats, with corresponding reductions in CSA and increases in MCA distensibility and incremental distensibility. These vascular effects suggest that PTUPB attenuates AD-associated hypertrophic remodeling, improves passive elastic properties of the vessel wall, and enhances vascular compliance in response to pressure changes. Collectively, these findings indicate that PTUPB restores cerebral autoregulation and vascular function in AD by normalizing vessel wall structure and mechanical properties.

The *ex vivo* vascular study was confirmed by transcriptomic profiling of primary VSMCs from AD animals, which showed that PTUPB significantly modulates pathways involved in contractility, extracellular matrix remodeling, inflammation, and oxidative stress. Pathway enrichment analysis revealed activation of focal adhesion, ECM– receptor interaction, PI3K–Akt, TGF-β, and VSMC contraction pathways (*Myh11, Mylk, Rhoa, Rock2*), suggesting improved VSMC tone, cytoskeletal anchoring, and reduced fibrosis.^76–80^ Enrichment of actin cytoskeleton, Rap1, and adherents junction pathways further pointed to enhanced contractile integrity and force transmission. In addition, modulation of MAPK, AGE–RAGE, TNF, shear stress/atherosclerosis, mTOR, and HIF-1 signaling suggested broad regulation of inflammation, oxidative stress, and metabolism.^81–84^ Together, these findings indicate that PTUPB acts on key pathways underlying VSMC dysfunction, providing a transcriptional explanation for the vascular improvements seen *in vivo*.

Consistent with the chronic study, acute vehicle-treated MCAs from AD rats exhibited impaired myogenic responses compared with vehicle-treated F344 vessels. Interestingly, low concentrations of PTUPB (0.1 and 1 µM) did not affect myogenic responses in F344 vessels but markedly inhibited the response at a higher concentration (10 µM). In contrast, PTUPB restored the myogenic response of AD MCAs at low concentrations (0.1 and 1 µM). This may reflect an anti-inflammatory action through sEH blockade and increased EET levels, since the IC₅₀ of PTUPB for sEH inhibition is 0.9 nM, compared with 1.3 µM for COX-2.^28^ At higher concentrations, however, PTUPB inhibited myogenic tone in AD vessels, similar to its effects in control F344 MCAs. This may be due to an additional elevation in EET levels as COX2 avidly metabolizes EETs to EHETs, especially when sEH activity is impaired.^85^

This dose-dependent response provided the basis to examine whether a similar concentration-dependent effect of PTUPB could be observed *in vivo* during functional hyperemia. Consistent with our previous work, functional hyperemic responses to whisker stimulation were markedly impaired in AD rats compared with F344 controls. At 1 μM, PTUPB markedly enhanced functional hyperemia in both strains relative to vehicle treatment. In F344 controls, this enhancement likely reflects sEH inhibition, which promotes the accumulation of vasodilatory EETs and reduces vasoconstrictor eicosanoid production, with additional elevations of EETs resulting from concurrent COX-2 inhibition. A similar effect was observed in AD rats; however, despite elevated sEH and COX-2 activity in AD, the improvement by PTUPB was not greater than in controls. This is likely due to additional factors contributing to neurovascular coupling, including PGs, EETs, NO, Kir2.1 channels in endothelial cells, and KATP channels in pericytes. Our recent finding that Kir2.1 expression is reduced in AD rats provides a plausible explanation for the attenuated hyperemic response despite PTUPB treatment.

In summary, the present study provides new evidence that dual inhibition of sEH and COX-2 improves cognition in AD, likely by enhancing myogenic responses and increasing cerebral artery distensibility. The cumulative concentration protocol used in this study minimized inter-vessel variability and allowed for within-vessel assessment of dose-dependent effects, but it also carries limitations. Although vessels were washed and re-equilibrated at baseline pressure between doses, residual drug or incomplete clearance cannot be ruled out. Similar considerations apply to the acute PTUPB treatment in the functional hyperemia study. Although we verified that the small molecular weight dye Evans blue (MW 961 Da) penetrated the thinned cranial window and dura to stain the brain surface, and that PTUPB (MW 543 Da) was applied topically for 10 minutes prior to measuring the hyperemic response, variability in bone thickness could still influence drug penetration, even though intracranial pressure was maintained. Finally, no direct measurements of PTUPB concentrations or sEH/COX-2 metabolites were performed, leaving uncertainty about the extent of target engagement under these conditions.

In conclusion, PTUPB alleviated cognitive decline in AD rats by restoring myogenic responses, improving vascular compliance, and reducing pathological remodeling, supporting its potential as a therapeutic strategy for AD.

## DECLARATIONS

### Author Contributions

F.F. conceived and designed the research; G.C.M., A.G., C.T., S.H.H., C.C., D.B., C.P., A.N., and P.O. performed the experiments; G.C.M., A.G., S.H.H., J.J.B., J.X., Y.L., S.B., T.J.L., A.P., and F.F. analyzed the data; G.C.M., A.G., S.H.H., T.J.L., K.M.W., C.M., P.O’H., Z.B., J.A.F., H.Y., B.D.H., R.J.R., and F.F. interpreted the results; G.C.M., A.G., J.X., Y.L., and F.F. prepared the figures; F.F. drafted the manuscript; all authors edited and approved the final version of the manuscript.

